# Intestinal catabolism of dietary fructose promotes obesity and insulin resistance via ileal lacteal remodeling

**DOI:** 10.1101/2025.08.18.670963

**Authors:** Miranda L. Lopez, Taekyung Kang, Ana Espeleta, Varvara I. Rubtsova, Jongwon Baek, Jakob Songcuan, Elena M. Moyer, Joohwan Kim, Won-Suk Song, Sunhee Jung, Nicholas D’Sa, Alexis Anica, Elise Tran, Yujin Chun, Wonsuk Choi, Ki-Hong Jang, Miranda E. Kelly, Ian J. Tamburini, Yasmine H. Alam, Johnny Le, Cuauhtemoc B. Ramirez, Raghu P. Kataru, Seon Pyo Hong, Dequina A. Nicholas, Katherine S. Xue, Gina Lee, Hosung Bae, Cholsoon Jang

## Abstract

High-fructose corn syrup (HFCS) consumption is a risk factor for obesity and metabolic syndrome, yet the underlying mechanisms are incompletely understood. Catabolism of dietary fructose primarily occurs in the small intestine and liver, with fructose breakdown in the liver being pathological, while small intestinal fructose clearance protects the liver. Here, we unexpectedly found that inhibition of fructose catabolism specifically in the small intestine mitigates fructose-induced obesity and insulin resistance. Mechanistically, blocking intestinal fructose catabolism reduces dietary fat absorption, which is associated with a decrease in the surface area of the ileal lacteals and alterations in gut microbiome. Fecal transplantation experiments revealed that such a microbiome stimulates the intestine-resident macrophages, promoting lacteal growth and boosting dietary fat absorption. Given the preclinical and clinical studies reporting the effect of fructose catabolism suppression on mitigating diet-induced obesity, our data suggest that such effects are partly mediated by intestinal lacteal remodeling.

**Significance Statement:** Here, we uncover a previously unappreciated link between intestinal fructose catabolism and ileal lacteal remodeling, suggesting the mechanisms by which fructose intake promotes obesity. Using mice lacking the fructose-processing enzyme specifically in the intestine, we show that blocking intestinal fructose metabolism protects against diet-induced obesity by reducing fat absorption. Changes in gut microbiome and immune cell interactions drive this effect.

## Introduction

Over the past century, sugar has steadily crept into modern diets — a trend largely attributed to the food industry’s embrace of HFCS to enhance the flavor of almost all processed food products. While dietary fat used to be regarded as the leading culprit of public health, in recent years, researchers have found that the added sugars in the diet, mainly in the form of fructose, are primarily contributing to alarmingly increased prevalence of obesity, diabetes, and metabolic dysfunction-associated steatotic liver disease (MASLD) (1–4). Although fructose and glucose, the two principal dietary sugars, carry the same amount of calories, their impact on the body differs drastically due to their distinct metabolism. For example, compared to glucose, fructose that reaches the liver acts as a potent inducer of *de novo* lipogenesis, leading to hepatic fat accumulation and increased adiposity across organs (5).

We and others previously demonstrated that fructose metabolism initiates in the small intestine (6–9), driven by the high expression of Ketohexokinase-C (*Khk-C*) (10). Genetic deletion of Khk-C in the small intestine under 10% sucrose drinking conditions causes fructose spillover to the liver, leading to excessive hepatic fructose catabolism and MASLD development (7). Conversely, individuals with essential fructosuria, a hereditary whole-body deficiency of *Khk*, rarely develop metabolic disorders not only because they avoid fructose intake due to fructose intolerance but also because ingested fructose is barely catabolized in the liver (11, 12). Instead, fructose is metabolized by hexokinase in other tissues (9, 13), with 10 ∼ 20% being excreted in the urine (14, 15).

Unlike glycolytic enzymes, *Khk* is not regulated allosterically (16). Accordingly, fructose catabolism occurs quickly once tissues encounter fructose. However, factors other than just enzyme expression can also influence fructose metabolism. For example, a high-fat diet-induced increase in villus length of the small intestine (17) may enhance fructose absorption and catabolism (17). However, the relationship between fructose metabolism and the complex substructures of the intestinal villi remains unknown.

Here, we found that mice lacking fructose catabolism, specifically in the small intestine, are protected against fructose-induced obesity. Unexpectedly, this phenotype was associated with decreased ileal lacteal surface area and poor dietary fat absorption. We further found the critical role of the gut microbiome and tissue-resident macrophages in regulating lacteal development, fat absorption, and adiposity. These data uncover a hitherto unrecognized connection between intestinal fructose catabolism, gut microbiome, and lacteal growth.

## Results

### Impact of intestinal *Khk-C* deletion on body fitness and glucose homeostasis

We have previously generated intestine-specific *Khk-C* deletion mice (*Khk-C*^ΔVilli^) (7) having *lox*P sites around a *Khk-C*-specific exon and intestine-specific Cre (*Villin*-Cre) (**Fig. 1A**). These mice exhibit intact *Khk-A* expression and a trend of decrease in other fructose-specific genes, such as *Aldolase B* (*AldoB*) and *Glucose transporter 5* (*Glut5*) (**Figs. 1B and S1A, B**). Impaired intestinal fructose clearance in these mice leads to excessive fructose spillover into the liver, resulting in increased hepatic lipogenesis and steatosis, even with a relatively low-dose fructose regimen (10% sucrose in drinking water) (7, 9). To expedite the pathogenic impact of fructose in mice, investigators often employed a high-dose (30%) *HFCS 55* in drinking water – composed of 45% glucose and 55% fructose (10, 18–20). Accordingly, in this study, we also adopted this regimen for studying the role of intestinal fructose catabolism in various metabolic pathophysiology other than hepatic steatosis.

**Figure 1.**
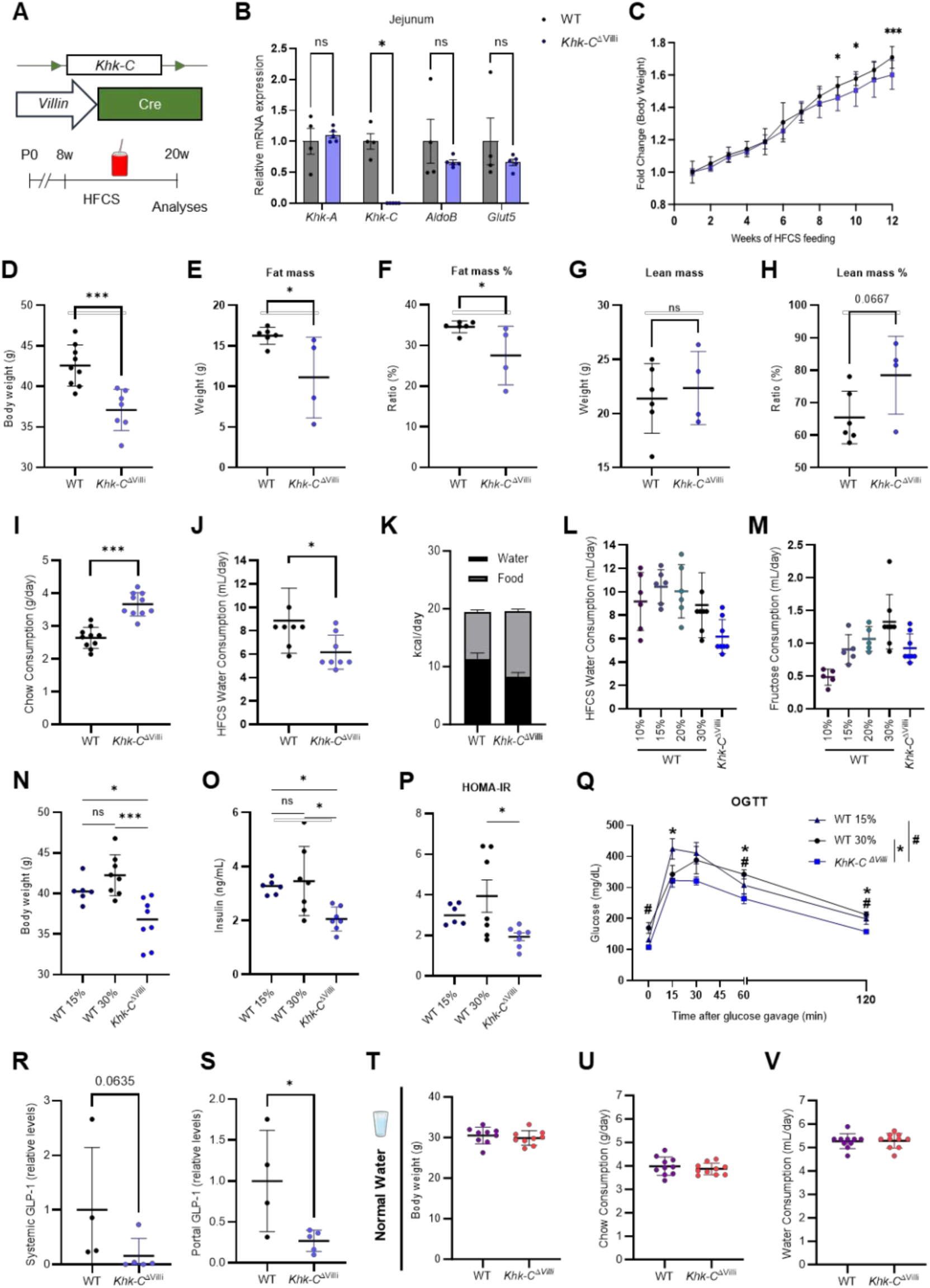
Suppression of intestinal fructose catabolism mitigates HFCS-induced obesity and improves glucose homeostasis. **(A)** 8-week-old WT or *Khk-C ^ΔVilli^* male mice were fed water containing 30% HFCS for 12 weeks. **(B)** Fructose catabolism gene expression in jejunum (n_WT_ = 4, n_Khk-C_ = 5). **(C, D)** Body weight changes and final body weight of mice on HFCS (n_WT_ = 9, n_Khk-C_ = 7). **(E, F)** Fat mass (**E**) and the ratio of fat mass to body weight (%) (**F**) (n_WT_ = 6, n_Khk-C_ = 4). **(G, H)** Lean mass (**G**) and the ratio of lean mass to body weight (%) (**H**) (n_WT_ = 6, n_Khk-C_ = 4). **(I-K)** Daily chow consumption (**I**), water intake (**J**), and total calorie intake (**K**) per mouse on HFCS (n = 10). **(L, M)** Daily water intake (**L**) and total fructose consumption (**M**) per mouse on varying concentrations of HFCS (n_WT10∼20%_ = 6, n_WT30%, Khk-C_ = 8). **(N)** Final body weight of mice on 15% or 30% of HFCS (n_WT15%_ = 6, n_WT30%, Khk-C_ = 8). **(O-Q)** Fasting insulin (**O**), homeostatic model assessment of insulin resistance (HOMA-IR) (**P**), and oral glucose tolerance test (OGTT) (**Q**) of mice on 15% or 30% of HFCS (n_WT15%_ = 6, n_WT30%, Khk-C_ = 4). **(R, S)** Systemic and portal blood levels of GLP-1 (n_WT_ = 4, n_Khk-C_ = 5). **(T)** Final body weight of mice on normal water (n = 9) **(U, V)** Daily chow (**U**) or water (**V**) consumption per mouse on normal water or HFCS (n = 10). Data are mean±s.d. ns, not significant. *p<0.05, **p<0.01, ***p<0.001 by one-way ANOVA followed by Tukey’s test (**N-Q**) or Student’s t-test.

After feeding 12 weeks of 30% HFCS water, we observed that *Khk-C*^ΔVilli^ mice were modestly yet significantly leaner than the littermate control (WT) mice (**Fig. 1C, D**). This was attributed to lower fat mass (**Fig. 1E, F**). Lean mass was not significantly different (**Fig.1G, H**). Consistent with our previous observation in *Khk-C*^ΔVilli^ mice under 10% sucrose feeding condition (7), *Khk-C*^ΔVilli^ mice showed higher chow consumption and lower water consumption than WT mice (**Fig. 1I, J**). These changes balanced out, leading to no difference in total caloric intake (**Fig. 1K**). Nevertheless, to eliminate potential influence of different fructose consumption on the body weight gain, we tested varying doses (10%, 15%, and 20%) of HFCS water in WT mice and compared 30% HFCS water consumption in *Khk-C*^ΔVilli^ mice (**Fig. 1L**). We found that providing 15% HFCS water to WT mice matched the fructose intake of *Khk-C*^ΔVilli^ mice fed with 30% HFCS (**Fig. 1M**).

Even after matching the total fructose consumption, we observed that *Khk-C*^ΔVilli^ mice still displayed lower body weight (**Fig. 1N**). Furthermore, *Khk-C*^ΔVilli^ mice showed significantly lower fasting insulin levels (**Fig. 1O**), although fasting glucose levels were similar (**Fig. S1C**). Calculation of Homeostatic Model Assessment of Insulin Resistance (HOMA-IR) and oral glucose tolerance test (OGTT) revealed that *Khk-C*^ΔVilli^ mice have improved glucose homeostasis compared to WT mice (**Fig. 1P, Q**).

To examine whether hormonal changes drive the observed metabolic improvements in *Khk-C*^ΔVilli^ mice, we conducted a Luminex-based quantitative screen for various incretins, adipokines, and pancreatic hormones in both the systemic and portal circulation (**Fig. S1D, E**). Interestingly, glucagon-like peptide 1 (GLP-1), the incretin that inhibits appetite, was decreased in the blood of *Khk-C*^ΔVilli^ mice (**Fig. 1R, S**), possibly explaining the increased chow intake of *Khk-C*^ΔVilli^ mice (**Fig. 1I**). These data suggest that, while GLP-1 receptor agonists have been successfully used to treat obesity (21), the reduced body weight in *Khk-C*^ΔVilli^ mice is not likely due to GLP-1 activation but rather due to other intrinsic mechanisms associated with intestinal fructose catabolism.

It was previously suggested that the endogenously produced fructose from glucose through the sorbitol pathway plays some roles in metabolic dysfunctions (22, 23). Consistent with this notion, the small intestine expresses *Khk-C* basally even in the absence of dietary fructose intake (24). We thus assessed the impact of intestinal *Khk-C* ablation on the body weight gain under the conditions of no dietary fructose either in chow or water. In contrast to HFCS-feeding conditions, *Khk-C*^ΔVilli^ mice showed no differences in body weight gain, food intake, or water intake compared to WT mice (**Fig. 1S-U**). Thus, intestinal *Khk-C* deficiency improves body fitness and glucose homeostasis only under the conditions of excessive dietary fructose intake.

### Impact of intestinal *Khk-C* deletion on dietary fat absorption

We were curious about the mechanism by which intestinal *Khk-C* ablation suppresses body weight gain during high-dose HFCS feeding. Given that *Khk-C*^ΔVilli^ mice showed lower body weight despite higher food consumption (**Fig. 1D, I**), we reasoned that their absorption of high-calorie nutrients such as fat may be negatively affected. Thus, we conducted a dietary lipid absorption assay; oral administration of soybean oil followed by the time-course measurements of circulating triglyceride (TG) levels (**Fig. 2A, B**). After 4 weeks of HFCS consumption, we did not observe any difference between *Khk-C*^ΔVilli^ and WT mice in circulating TG levels after soybean oil provision (**Fig. S2A, B**). Strikingly, after 8 weeks of HFCS consumption, *Khk-C*^ΔVilli^ mice started to exhibit a significant reduction in circulating TG levels (**Fig. S2C, D**). Furthermore, the phenotype became even more prominent after 12 weeks of HFCS feeding (**Fig. 2C, D**), suggesting the importance of chronic fructose exposure. To verify the difference in dietary lipid absorption in an alternative way, we measured TG contents in the feces. *Khk-C*^ΔVilli^ mice showed substantially higher TG content in the feces than WT mice (**Fig. 2E**), further implicating the reduced dietary fat absorption in *Khk-C*^ΔVilli^ mice. Thus, we concluded that ablating intestinal fructose catabolism impairs dietary fat absorption after long-term HFCS consumption.

**Figure 2.**
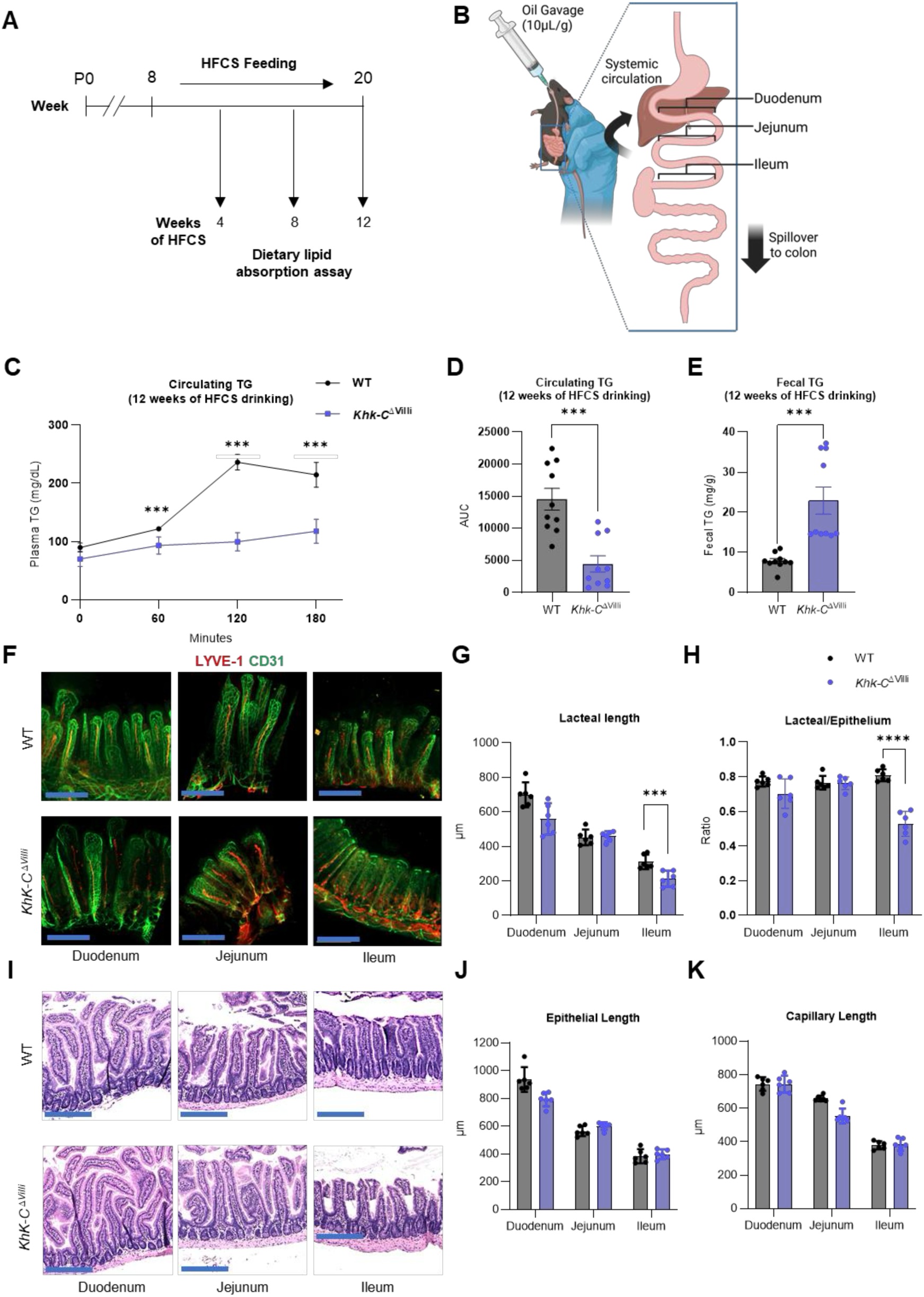
Suppression of intestinal fructose catabolism reduces dietary fat absorption and lacteal shortening under HFCS consumption. **(A, B)** During 12-week HFCS feeding, dietary fat absorption assay was performed on weeks 4, 8, 12 of HFCS feeding, using oral soybean oil provision followed by circulating and fecal triglyceride (TG) measurements. **(C)** Measurements of circulating TG levels **(C)** and the corresponding area under curve (AUC) **(D)** after 12 weeks of HFCS drinking (n = 10). **(E)** Measurements of TG excretion in feces collected 120 minutes post-oil gavage after 12 weeks of HFCS drinking (n = 10). **(F)** Representative immunofluorescence images of intestinal whole mounts showing LYVE-1-positive lacteals (red) and CD31-positive capillaries (green) in mice on HFCS feeding for 12 weeks. Scale bars, 200 μm. **(G, H)** Lacteal length (**G**) and the ratio of lacteal/epithelial length (**H**) in intestinal tissues of mice on HFCS feeding (n = 6). **(I)** Representative H&E images of intestinal tissues of mice on HFCS feeding. Scale bars, 200 μm. **(J, K)** Epithelial (**J**) and capillary length (**K**) of intestinal tissues in mice on HFCS feeding (n = 6). Data are mean±s.d. ***p<0.001, ****p<0.0001 by Student’s t-test.

### Impact of intestinal *Khk-C* deletion on lacteal growth

Next, we investigated the mechanisms of how *Khk-C* deficiency in the small intestine decreases dietary fat absorption. Fat transport in the small intestine is initiated by the uptake of dietary fatty acids into enterocytes via various transport proteins (Cd36, Fatp1, Fatp4, Fabp2), followed by reassembly into TGs (25). Such TGs are then transported through intestinal lacteals, the small lymphatic vessels within intestinal villi (26). We found no significant changes in mRNA expression of major fatty acid transport proteins across the duodenum, jejunum, and ileum of *Khk-C*^ΔVilli^ mice (**Fig. S2E-G**). Next, we assessed the lacteal length, which is also critical for lipid absorption (27, 28). Whole-mount immunostaining revealed that, while lacteal length was similar between *Khk-C*^ΔVilli^ and WT mice in the duodenum and jejunum, lacteal length in *Khk-C*^ΔVilli^ was significantly shorter than WT mice, specifically in the ileum (**Fig. 2F-H**). In contrast, we did not observe any difference in epithelial or capillary length between *Khk-C*^ΔVilli^ and WT mice (**Fig. 2I-K**). Moreover, under normal water drinking conditions without dietary fructose, neither intestinal lacteal length nor epithelial/capillary length showed apparent change in *Khk-C*^ΔVilli^ mice (**Fig. S2H-M**). Thus, excessive fructose intake triggers lacteal shortening in the ileum, which likely results in the decrease in dietary fat absorption in animals lacking intestinal fructose catabolism.

### Impact of intestinal *Khk-C* deletion on gut microbiota and villus macrophages

We next sought to identify the mechanism that underlies the decreased ileal lacteal length in *Khk-C*^ΔVilli^ mice. Previous research has identified the gut microbiota as a crucial regulator of lacteal growth by affecting the release of growth factors from intestinal villus macrophages (27). We thus hypothesized that, blunted fructose catabolism by the host intestine changes nutritional environment of gut microbiota, affecting their ecosystem fructose to be retained within the intestinal lumen, where microbiome can consume more fructose and change compositions/metabolism, eventually affecting the lacteal length. To investigate this idea, we first conducted 16S rRNA sequencing of gut microbiota at multiple time frames – 0, 1, 3, 7, 14, 28 and 84 days after HFCS feeding (**Fig. 3A**).

**Figure 3.**
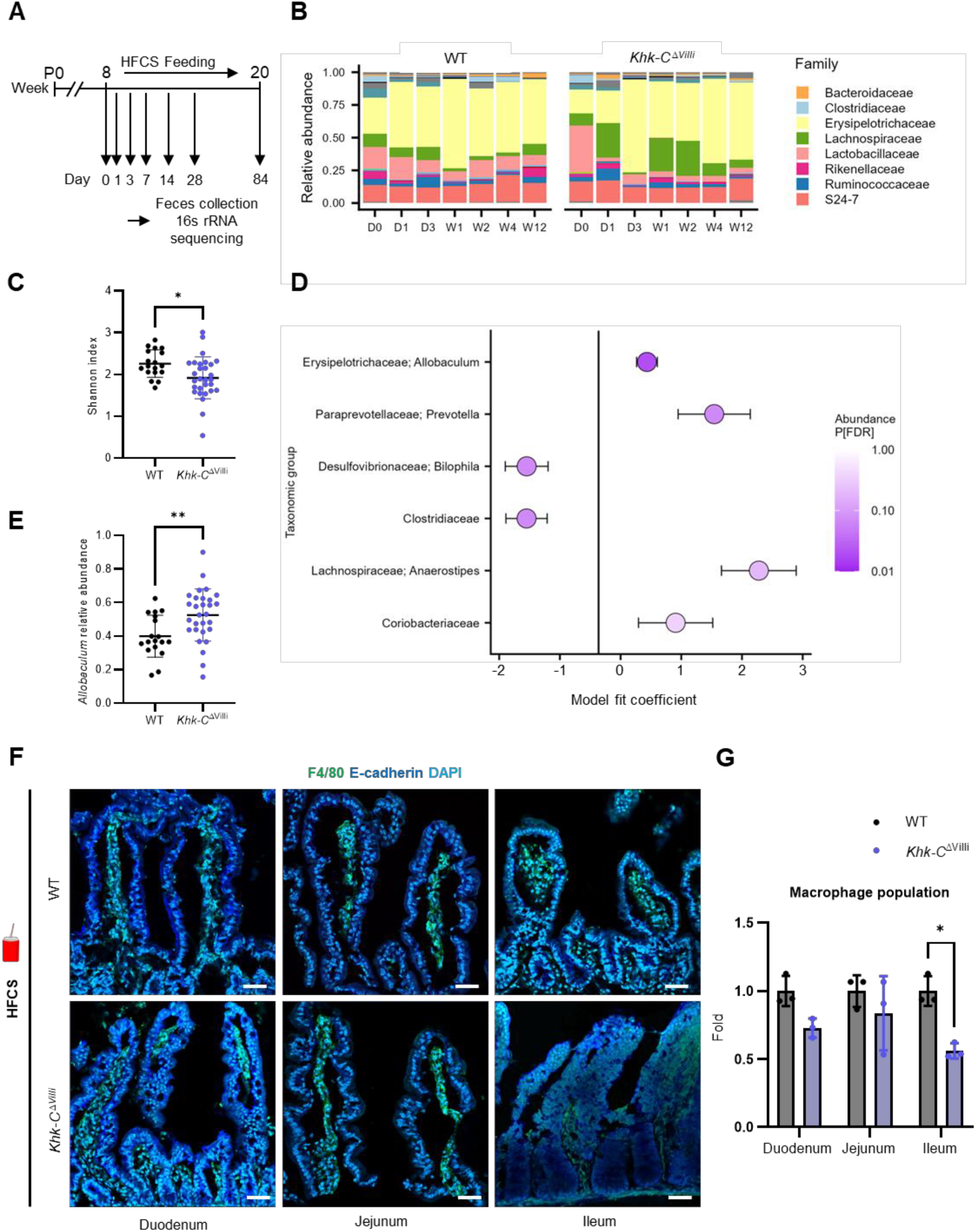
Suppression of intestinal fructose catabolism alters gut microbiome diversity and reduces villus macrophages under HFCS consumption. **(A)** Schematic overview of fecal 16S rRNA sequencing across multiple days of HFCS intake. **(B)** Representative bacterial family compositions from a WT and *Khk-C ^ΔVilli^* mouse at each timepoint. **(C)** Alpha diversity (Shannon index) across all samples for WT and *Khk-C ^ΔVilli^* mice post-day 14 of HFCS intake (n_WT_ = 18, n_Khk-C_ = 28). **(D)** Differential abundance results from MaAsLin3 showing the five taxa with the strongest associations between WT and *Khk-C ^ΔVilli^* mice post-day 14 of HFCS intake n_WT_ = 18, n_Khk-C_ = 28). X-axis represents the model fit coefficient (β) from MaAsLin3, indicating the direction and magnitude of association with the *Khk-C ^ΔVilli^*genotype (positive = enriched in *Khk-C ^ΔVilli^*, negative = enriched in WT). Only *Allobaculum spp*. was significant with a false discovery rate <0.05. **(E)** Relative abundance of Allobaculum in WT and *Khk-C ^ΔVilli^*mice post-day 14 of HFCS intake (n_WT_ = 18, n_Khk-C_ = 28). **(F)** Representative immunofluorescence images of intestinal cryosections showing F4/80-positive macrophages (green), E-cadherin-positive epithelial cells (blue), and nuclei (DAPI, turquoise). Scale bars, 50 μm. **(G)** Relative abundance of macrophages in mice on HFCS for 12 weeks (n = 3). Data are mean±s.d. *p<0.05, **p<0.01 by Student’s t-test.

Following HFCS feeding, we observed significant decreases in microbiome diversity in *Khk-C*^ΔVilli^ mice compared to the microbiota in WT mice (**Fig. 3B, C**), a trait commonly seen in patients with metabolic disorders (29). Notably, MaAsLin3 differential abundance analysis revealed that *Allobaculum spp.* (family Erysipelotrichaceae) became a significantly more abundant bacterial species in *Khk-C*^ΔVilli^ mice after at least two weeks of HFCS feeding (**Fig. 3D, E**). This finding was intriguing because prior studies have shown that members of the Erysipelotrichaceae family expand in mice on high-fat and high-sugar diets, affecting host lipid metabolism (30, 31).

Considering that Erysipelotrichaceae directly influences the intestinal immune system (32) and that villus-resident macrophages are critical sources of growth factors for lacteal growth (27, 33), we next examined villus macrophages. F4/80 immunostaining revealed that *Khk-C*^ΔVilli^ mice have a markedly reduced number of macrophages in the ileum but not in the duodenum or jejunum, compared to WT mice (**Fig. 3F, G**). However, *Khk-C*^ΔVilli^ mice fed normal water without HFCS did not show any change in the number of F4/80-positive macrophages across intestinal segments (**Fig. S3A, B**). These results suggest that microbiome alterations affect villus-resident macrophages in HFCS-fed *Khk-C*^ΔVilli^ mice, contributing to the shortening of lacteals.

### The role of microbiota in the ileal lacteal growth and fat absorption

To directly test whether the microbiota is a key contributing factor to ileal lacteal growth and dietary fat absorption, we conducted fecal transplant experiments. To this end, we generated two groups of fecal donor mice: *Khk-C*^ΔVilli^ and WT mice fed 30% HFCS for 12 weeks (**Fig. 4A**). In parallel, the recipient mice (WT) were first treated with a cocktail of antibiotics for a week to eliminate the gut microbiota. Then, we transplanted freshly isolated fecal contents from the donor mice to the recipient mice twice a week orally while feeding the recipient mice 30% HFCS for 8 weeks. Encouragingly, the mice that received fecal contents from *Khk-C*^ΔVilli^ donors exhibited significantly lower body weight gain, compared to the mice that received fecal contents from WT donors (**Fig. 4B**).

**Figure 4.**
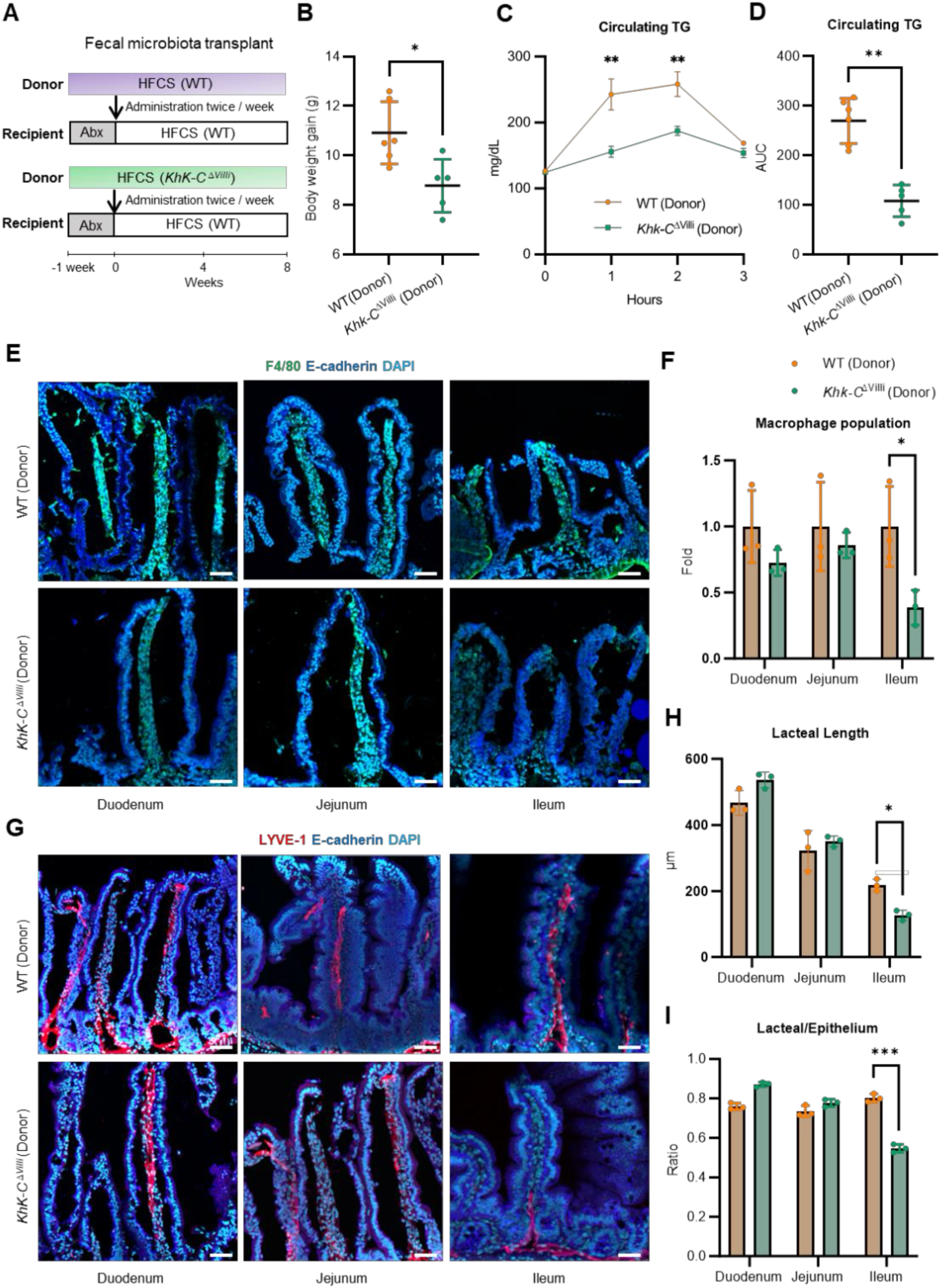
Fecal microbiota transplant phenocopies intestine-specific *Khk-C* depletion mice under HFCS consumption. **(A)** Schematic of fecal transplant (twice/week) from donors (WT or *Khk-C ^ΔVilli^* mice on HFCS for 12 weeks; n = 6) to recipients (WT after antibiotics) with HFCS drinking for 8 weeks. **(B)** Body weight gain of the recipient mice after 8 weeks of fecal transplant and HFCS drinking (n_WT(Donor)_ = 6, n_Khk-C(Donor)_ = 5). **(C, D)** Measurements of circulating TG levels (**C**) and the AUC (**D**) in the recipient mice following soybean oil gavage (n_WT(Donor)_ = 6, n_Khk-C(Donor)_ = 5). **(E)** Representative immunofluorescence images of intestinal cryosections showing F4/80-positive macrophages (green), E-cadherin-positive epithelial cells (blue), and nuclei (DAPI, turquoise) in the recipient mice after 8 weeks of fecal transplant and HFCS drinking. Scale bars, 50 μm. **(F)** Relative abundance of macrophages in the recipient mice (n = 3). **(G)** Representative immunofluorescence images of cryosections showing LYVE-1-positive lacteals (red), E-cadherin-positive epithelial cells (blue), and nuclei (DAPI, light blue) in the recipient mice after 8 weeks of fecal transplant and HFCS drinking. Scale bars, 50 μm. **(H, I)** Lacteal length (**H**) and the ratio of lacteal/epithelial length (**I**) in intestinal tissues of the recipient mice after 8 weeks of fecal transplant and HFCS drinking (n = 3). Data are mean±s.d. *p<0.05, **p<0.01, ***p<0.001 by Student’s t-test.

Motivated by this observation, we performed the oral lipid absorption assay. We found reduced dietary fat absorption in the recipient mice that received fecal contents from *Khk-C*^ΔVilli^ donors compared to the mice that received fecal contents from WT mice (**Fig. 4C, D**). These recipient mice from *Khk-C*^ΔVilli^ donors also displayed a significantly decreased number of villus macrophages in the ileum, but much less pronounced effects in the duodenum or jejunum (**Fig. 4E, F**), recapitulating the findings in the donor mice (**Fig. 3F, G**).

Finally, whole-mount immunostaining revealed that the recipient mice from *Khk-C*^ΔVilli^ donors exhibited a reduction in lacteal length as well as the ratio of lacteal to epithelial length in the ileum (**Fig. 4G-I**). However, we did not observe any difference in epithelial or capillary length (**Fig. S4A, B**), again recapitulating the findings in the donor mice (**Fig. 2F-H**). Altogether, these data suggest that the gut microbiota act as a critical mediator of the effects of impaired intestinal fructose catabolism in remodeling ileal macrophage populations, lacteal growth, and ultimately, dietary fat absorption, under the conditions of high-dose HFCS consumption. These processes likely contribute to the observed reduction in body weight gain and improved glucose homeostasis in mice lacking intestinal fructose catabolism.

## Discussion

In a contemporary society where excessive fructose and calories are widespread in processed foods, the small intestine’s role in absorbing and processing dietary nutrients becomes even more critical in determining the trajectory between health and disease. While many studies have investigated the impact of high-fat diet consumption on the absorption of other nutrients (18, 35, 36), the effect of fructose consumption or its metabolic capacity on the absorption of other nutrients is relatively understudied. Here, we found an unexpected role of small intestinal fructose catabolism in modulating gut microbiome compositions, ileum-specific lacteal growth, dietary fat absorption, and eventually whole-body metabolic fitness following chronic consumption of HFCS.

Individuals having loss-of-function mutations in *Khk* have been known to be resistant to fructose-associated pathologies, including obesity, diabetes, and MASLD (11, 12). Without fructose consumption, such individuals have no overt phenotypes, consistent with our data in genetically modified mice. Accordingly, *Khk* inhibitors have been developed by pharmaceutical companies for treating metabolic disorders, with some notable success in clinical trials (14, 34–36). While these drugs likely exert systemic effects on all organs that express *Khk* (37), including the liver, intestines, kidneys and adipose tissues, only the liver has garnered attention (38). Our results suggest that the reported clinical benefits are at least partly attributed to the drug’s actions on the small intestine and its suppressive impact on dietary fat absorption.

Conversely, the appropriate absorption of dietary fat, especially essential fatty acids, is crucial not only for maintaining body weight but also for supporting organ health. Common intestinal pathologies—such as irritable bowel syndrome (42) or intestinal cancer (43) — negatively influence intestinal functions, including fructose catabolism, through chronic inflammation and epithelial cell de-differentiation. While ∼ 20% of cancer patients experience cancer-related cachexia, ∼ 60% of those with intestinal cancer suffer from this condition (39). However, no sufficient data are available regarding their fructose consumption or dietary fat absorption capacity. It will be necessary to investigate whether impaired dietary fat absorption contributes to cachexia progression and whether intravenous fat provision may help mitigate the disease in these patients (40).

Beyond the host-specific factors, our fecal transplant data suggest the importance of the gut microbiota in maintaining lacteal integrity, particularly in the ileum, under the condition of HFCS consumption. It is plausible to hypothesize that the gut microbiota play different roles in different compartments of the small intestine, a topic that is poorly understood. Only recently have studies begun to reveal that the microbiota in the ileum not only shows a closer resemblance to the overall gut microbiota than those in other small intestinal segments (41) but also reacts similarly to fructose feeding (42). Further studies will be necessary to better understand how HFCS consumption affects lacteal integrity selectively in the ileum through modulating gut microbiome and eventually the host’s dietary fat absorption.

In this regard, our findings that the gut microbiota likely influence the population of ileal macrophages to affect lacteal length under the condition of HFCS consumption present a previously unrecognized interaction between fructose and gut immune-microbe interactions. Although our sequencing data highlight the Erysipelotrichaceae family as a potentially important microbiota in this process, and members of the Erysipelotrichaceae family have been associated with diet-induced microbiome changes and lipid metabolism in obese mice (30–32, 43), the mechanism by which this effect is mediated remains unclear, particularly whether it is mediated by macrophage depletion. Looking forward, identifying specific microbiome species responsible for promoting villus macrophage populations and lacteal growth will be instrumental in developing new probiotics to therapeutically modulate dietary fat absorption.

## Materials and Methods

### Mouse studies

Animal studies followed protocols approved by the Institutional Animal Care and Use Committee of the University of California, Irvine. 8-week-old male C57BL/6 mice were purchased from Jackson Laboratory and group-housed (2-3 mice/cage) on a standard light-dark cycle (7:00– 19:00) with free access to chow (Teklad™ 2020X) and water. *Khk-C* floxed mice (7) were bred with *Villin*-Cre mice (stock no. 004586; Jackson Laboratory) to generate intestine-specific *Khk-C* knockout mice. For HFCS provision, mice were provided either ad libitum H_2_O or a 55% fructose : 45% glucose mixture in their drinking water. Daily water and food intake were determined by measuring the total consumption in a cage and dividing by the number of mice in the cage. For antibiotics treatment, a cocktail of antibiotics, including 1 g L^−1^ ampicillin, neomycin, metronidazole, and 0.5 g L^−1^ vancomycin, was dissolved in HFCS water. For gut bacteria transplantation, fecal pellets were freshly collected from donor mice, immediately dissolved in PBS (2 mL), centrifuged at maximum speed at 4°C for 10 minutes, and the supernatant was transferred to antibiotics-treated recipient mice via oral gavage (200 µL/mouse) with a plastic feeding tube (Instech Laboratories) every 3-4 days for 8 weeks. Body composition was assessed using EchoMRI™ Whole Body Composition Analyzer (Houston, TX, United States), which provides measurements of whole body fat and lean mass.

### Glucose tolerance test

For glucose tolerance test, mice received D-glucose in sterile saline at a dosage of 2g/kg via oral gavage after 12 hours of fasting. Glucose levels were determined from a tail snip with a hand-held glucometer (Accu-Check Guide) in the basal state and 15-, 30-, 60-, and 120-min following glucose administration. For insulin measurements, serum was collected after 6 hours of fasting. Insulin levels were measured using an Ultra Sensitive Mouse Insulin ELISA Kit (catalog no. 90080; Crystal Chem). The HOMA-IR was calculated with the following formula (glucose levels in mmol/L, insulin levels in uIU/mL): *HOMA-IR = (glucose x insulin) /* 22.5.

### Circulating hormone measurements

For circulating appetite and energy homeostasis-related hormone measurements, serum samples were centrifuged for 30 sec to remove debris before being applied to a 384-well plate for analysis by multiplexing bead-based ELISA using the MMHE-44K metabolic Milliplex Hormone kit (Millipore). Antibodies and magnetic beads were diluted 1:1 with assay buffer. 10 µL of antibodies, magnetic beads, and serum samples were utilized to adjust the manufacturer’s protocol to our 384-well plate format. Each sample, standard, and quality control were assayed in technical duplicates, with the mean fluorescence intensity (MFI) value used for analyses. Plates were washed in between incubations using a BioTek 406 Touch plate washer (BioTek) and read using the Luminex xMAP INTELLIFLEX system. Samples were diluted at 1:7 (sample: assay buffer) to allow abundant hormones to be measured in the linear range of the instrument. Data analysis was performed using Belysa immunoassay software.

### Oral lipid absorption assay

For the dietary lipid absorption assay, mice received soybean oil via oral gavage (10 µL g^−1^) after 6 hours of fasting. Blood was collected via tail snip to measure circulating triglyceride after different durations. Feces were snap-frozen in liquid nitrogen with a pre-cooled Wollenberger clamp. Multiple cohorts were used, and data were combined if there was no statistical difference between cohorts. Triglyceride levels were measured using a Triglyceride Colorimetric Assay Kit (catalog no. 10010303; Cayman Chemicals).

### Immunofluorescence imaging

For whole-mount microscopy of intestinal segments, we adapted a previously established protocol (44). Intestinal samples were prepared carefully by removing the intestine from the mesentery of the pyloric ring with forceps. All samples were kept in a 15-cm, 6-well culture plate filled with ice-cold phosphate-buffered saline (PBS). The extracted intestines were divided into three equal segments: proximal (duodenum), mid (jejunum), and distal (ileum), and feces and other luminal contents were flushed out using a syringe. The samples were longitudinally cut along the mesenteric border to expose the lumen and further sectioned into small pieces measuring 0.5-1 cm, then dissected into strips of intestinal villi. Tissue samples were fixed in 4% buffered paraformaldehyde overnight at 4°C. After fixation, tissues were sequentially dehydrated with 10% sucrose for 2 hours and 20% sucrose overnight at 4°C. Dehydrated tissues were subsequently frozen and embedded in Frozen Section Media (Leica) for cryosection and cut into 20-μm-thick sections using a Cryocut Microtome (Leica). Samples were then blocked in protein block serum (Agilent) with 0.3% Triton X-100 (Thermo Fisher) for 1 hour and incubated with primary antibodies in antibody diluent (Agilent) overnight at 4°C, followed by incubation with anti-mouse secondary antibodies for 2 h at room temperature. Tissue sections were mounted with Vectashield plus DAPI (4’,6-diamidino-2-phenylindole) (Vector Labs) and imaged using an LSM 980 confocal microscope (Zeiss). The following primary antibodies were used: anti-CD31 (rat monoclonal, clone MEC 13.3, BD Bioscience; anti-F4/80 (rat monoclonal, clone BM8, BioLegend); anti-E-cadherin (goat polyclonal, R&D); anti-LYVE-1 (rabbit polyclonal, AngioBio). Alexa 488-, Alexa 594-, and Alexa 647-conjugated secondary antibodies were purchased from Jackson ImmunoResearch. Morphometric analyses were conducted using data processed through LSM Zen (Carl Zeiss), ImageJ software (NIH), or Imaris (Bitplane). The blood capillary, lacteal, and epithelial lengths (villus heights) were measured for each villus. Given that lacteal length varies significantly but tends to correlate with villus or capillary length, we also calculated the ratios of lacteal to villus and lacteal to capillary lengths. These relative measurements are known to remain consistent along the entire length of the intestine (45). Each measurement represents the mean of six measurements per sample. Macrophage density was quantified by measuring the F4/80^+^ area across five 0.0265 mm² fields per sample, with the results presented as relative values normalized to the area.

### Histology

For hematoxylin and eosin (H&E) staining, freshly collected intestinal tissues were fixed in 4% paraformaldehyde overnight, embedded in paraffin, sectioned, and stained with H&E. Tissues were submitted to the Experimental Tissue Shared Resource Facility at the University of California Irvine. Images were captured using a high-resolution image scanner (Ventana DP200 ROCHE). For Oil Red O staining, freshly collected hepatic tissues were fixed in 4% paraformaldehyde overnight, dehydrated by titrating in sucrose (10%, 20%) and embedded in Frozen Section Media (Leica) for cryosection using a Cryocut Microtome (Leica), and stained with Oil Red O. Images were captured using a Nano-Zoomer Digital Pathology scanner (Hamamatsu, Japan).

### RT–qPCR

RNA samples were prepared using TRIzol® Reagent (Invitrogen) according to the manufacturer’s instructions. RNA was reverse transcribed to cDNA using the iScript kit (Bio-rad). The resulting cDNA was analyzed by quantitative RT-PCR (qPCR) using SYBR green master mix (Life Technologies) on QuantStudio6 Real-Time PCR system (Life Technologies). Relative mRNA expression was calculated from the comparative threshold cycle (Ct) values relative to housekeeping genes *Actin*, *36b4*, and *Tbp*. Primer sequences for RT–qPCR are: *Glut5* (forward: TCTTTGTGGT AGAGCTTTGGG, reverse: GACAATGACACAGACAATGCTG), *Khk-a* (forward: TTGCCGATTTT GTCCTGGAT, reverse: CCTCGGTCTGAAGGACCACAT), *Khk-c* (forward: TGGCAGAGCCAGG GAGAT, reverse: ATCTGGCAGGTTCGTGTCGTA), *Aldob* (forward: CACCGATTTCCAGCCCTC, reverse: GTTCTCCACCTTTATCCTTTGC), *Acly* (forward: CAGCCAAGGCAATTTCAGAGC, rev erse: CTCGACGTTTGATTAACTGGTCT), *Acss2* (forward: ATGGGCGGAATGGTCTCTTTC, rev erse: TGGGGACCTTGTCTTCATCAT), *Fasn* (forward: GGAGGTGGTGATAGCCGGTAT, reverse: TGGGTAATCCATAGAGCCCAG), *Scd1* (forward: TTCAGAAACACATGCTGATCCTCATAATT CCC, reverse: ATTAAGCACCACAGCATATCGCAAGAAAGT), *Tbp* (forward: CCCTATCACTCC TGCCACACCAGC, reverse: GTGCAATGGTCTTTAGGTCAAGTTTAC), *Actin* (forward: CCCTGT ATGCTCTGGTCGTACCAC, reverse: GCCAGCCAGGTCCAGACGCAGGATG), *36b4* (forward: GGAGCCAGCGAGGCCACACTGCTG, reverse: CTGGCCACGTTGCGGACACCCTCC).

### 16S rRNA gene amplicon sequencing and analysis

Fresh feces samples were collected at 9 AM, snap-frozen, and stored at -80 °C for 16S rRNA sequencing. The fecal DNA was extracted using the ZymoBIOMICS 96 DNA Kit (Zymo Research). 16S rRNA amplicon PCR was performed targeting the V4-V5 region using the EMP primers (515F (barcoded) and 926R) (46, 47). The sequencing library was prepared using an innovative library preparation process, where PCR reactions were performed in real-time PCR machines to control cycles and thus limit PCR chimera formation. The library was sequenced on Illumina MiSeq v3 chemistry with a PE300 sequencing length. Unique amplicon sequence variants were inferred from raw reads using the DADA2 pipeline. The sequences were assigned a taxonomic classification using the August 2013 greengenes database (greengenes.secondgenome.com), trained with the primer pairs that were used to amplify the 16S region. The extracted region was truncated at 243 bp for single read analysis which should give the best taxonomical identification. Composition visualization, alpha-diversity, and beta-diversity analyses were performed using Qiime v2. Taxa with differential abundance and prevalence between the WT and *Khk-C*^ΔVilli^ groups were identified by running MaAsLin3 (48) using parameters --normalization=”TSS “ --transform=”LOG” --fixed_effects=(“*Khk-C*^ΔVilli^.gene”) -- reference=(“*Khk-C*^ΔVilli^.gene, WT”) on samples from days 14-84. *Allobaculum spp.* was the only genus with significant differential abundance between the WT and *Khk-C*^ΔVilli^ groups at an FDR threshold of 0.05, and no taxa had significantly different prevalence at this threshold.

### Statistical analysis

Otherwise indicated, a two-tailed, unpaired Student’s t-test was used to calculate P values, with P < 0.05 used to determine statistical significance. One-way ANOVA followed by Tukey’s test was used for multiple comparisons. Outliers were determined by 1.5 times the interquartile range (1.5 * IQR below Q1 or 1.5 * IQR above Q3) (49, 50)

## Author Contributions

H.B. and C.J. conceived and supervised the study. M.L.L. and T.K. performed most experiments with the help of V.I.R., J.B., J.S., and E.M.M. M.L.L. performed multiplex hormone assays with supervision from D.A.N. H.B. and S.J. performed 16S rRNA gene amplicon sequencing, and A.E. and W-S.S. analyzed the data with supervision from K.S.X. J.K. and N.D. performed histological analysis with supervision from G.L. A.A. and E.T. provided technical assistance. Y.C., W.C., K-H.J., M.E.K., I.J.T., Y.H.A., J.L., and C.B.R. contributed intellectually. R.P.K. and S.P.H. provided immune and lymphatic expertise. H.B. and C.J. wrote the manuscript with comments from all authors.

## Classification

Biological Sciences / Physiology

## Acknowledgments

We dedicate this manuscript to our beloved colleague, mentor, and friend, Dr. Gina Lee, who recently lost her battle to breast cancer. She remains forever in our hearts. This work was supported by the National Research Foundation of Korea (2021R1A6A3A14039132, H.B; 2021R1A6A3A-14039681, S.J), Pew Foundation (C.J.), and National Institutes of Health grants (R01-AA029124, R21-AA030358 to C.J). We thank the UCI microbiome center for providing 16S rRNA sequencing. Illustrations were created with BioRender.com.

## SUPPLEMENTARY FIGURE LEGENDS

**Supplementary Fig 1.**
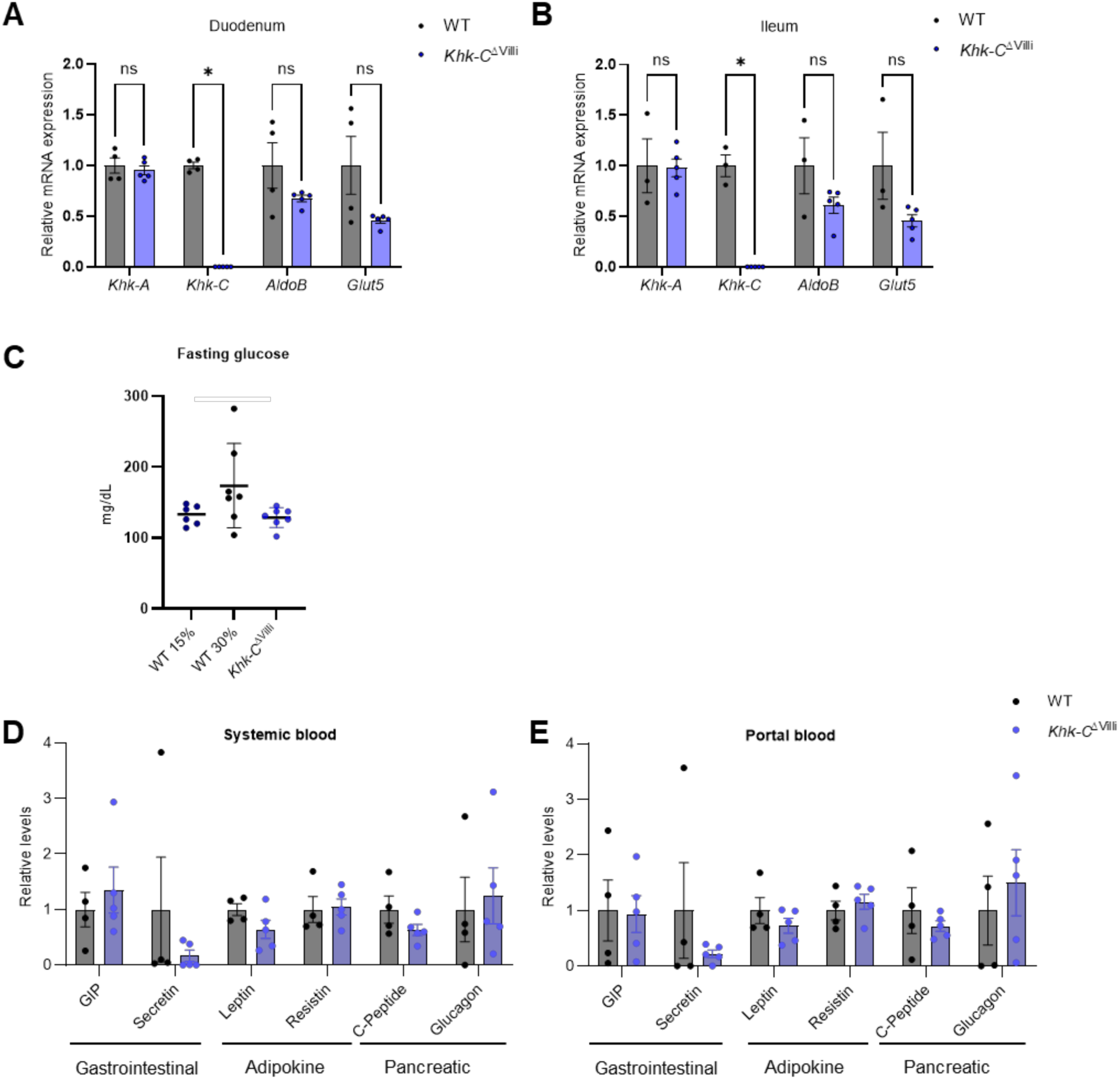
Suppression of intestinal fructose does not affect body weight under normal water conditions. **(A, B)** Fructose catabolism gene expression in duodenum and ileum (n_WT_ = 4, n_Khk-C_ = 5). **(C)** Fasting glucose levels (n_WT15%_ = 6, n_WT30%, Khk-C_ = 7) . **(D, E)** Systemic and portal blood levels of hormones (n_WT_ = 4, n_Khk-C_ = 5). Data are mean±s.d. ns, not significant. *p<0.05, **p<0.01 by Student’s t-test.

**Supplementary Fig 2.**
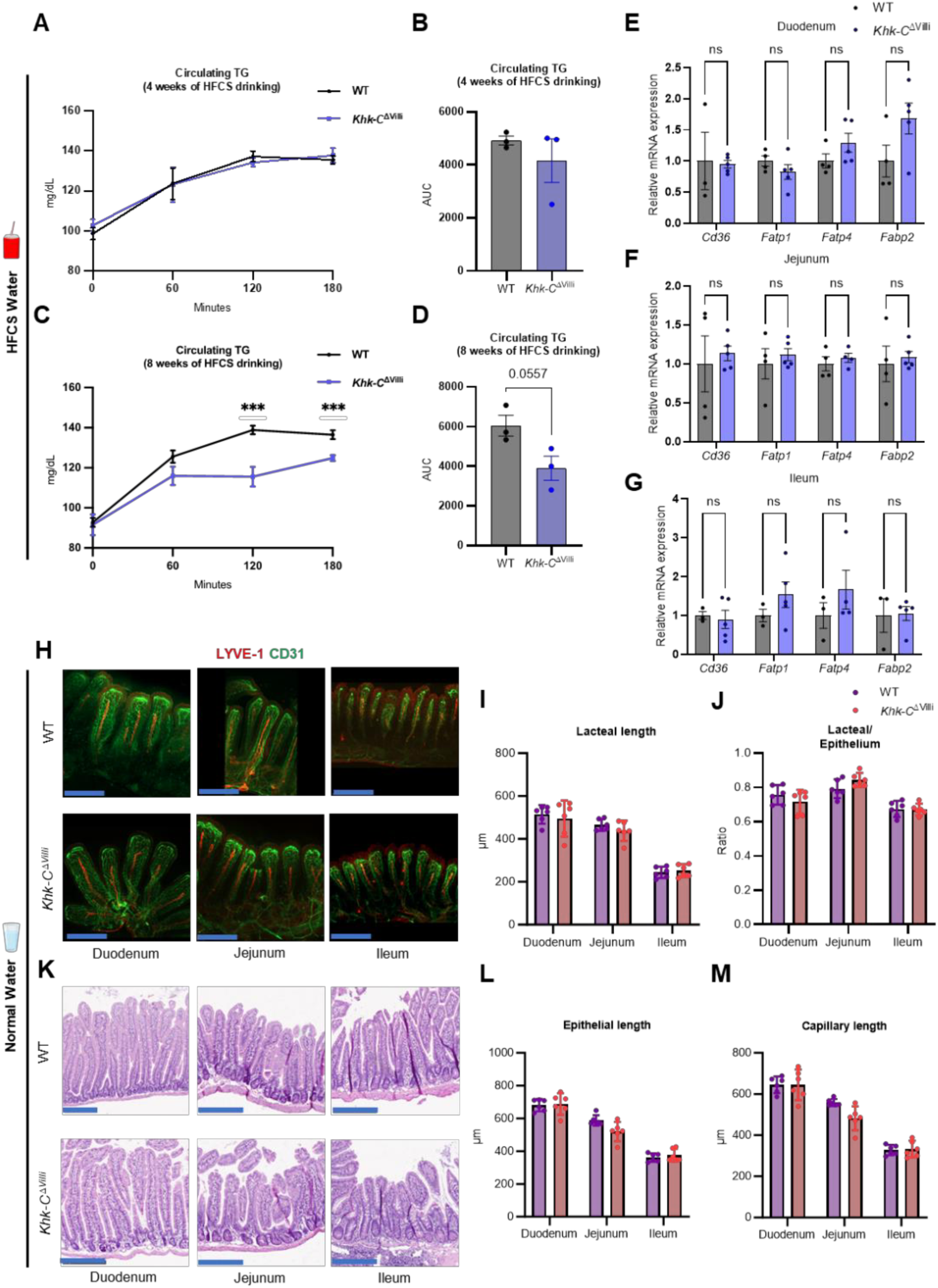
Suppression of intestinal fructose catabolism reduces dietary fat absorption without altering fatty acid transporter expression under HFCS consumption. **(A, B)** Measurements of circulating TG levels **(A)** and the corresponding area under curve (AUC) **(B)** after 4 weeks of HFCS drinking (n = 3). **(C, D)** Measurements of circulating TG levels **(C)** and the AUC **(D)** after 8 weeks of HFCS drinking (n = 3). **(E-G)** Relative fatty acid transporter gene expression in duodenum, jejunum, and ileum (n_WT_ = 4, n_Khk-C_ = 5). **(H)** Representative immunofluorescence images of intestinal whole mounts showing LYVE-1-positive lacteals (red) and CD31-positive capillaries (green) in mice on normal water feeding for 12 weeks. Scale bars, 200 μm. **(I, J)** Lacteal length (**I**) and the ratio of lacteal/epithelial length (**J**) in intestinal tissues of mice on normal water feeding (n = 6). **(K)** Representative H&E images of intestinal tissues of mice on normal water feeding. Scale bars, 200 μm. **(L, M)** Epithelial (**L**) and capillary length (**M**) of intestinal tissues in mice on normal water feeding (n = 6). Data are mean±s.d. ns, not significant. ***p<0.001 by Student’s t-test.

**Supplementary Fig 3.**
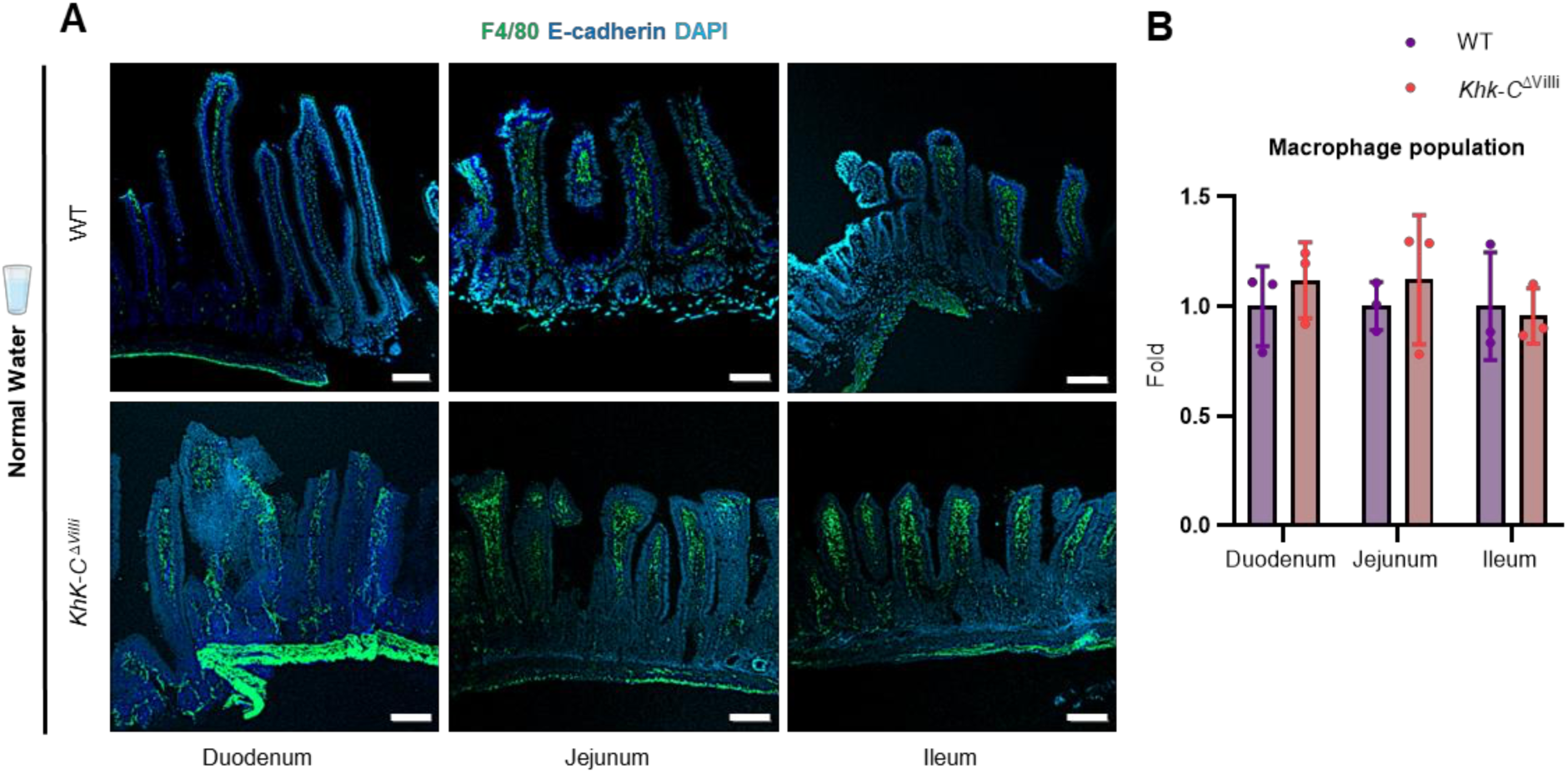
Suppression of intestinal fructose catabolism does not reduce villus macrophage population under normal water conditions. **(A)** Representative immunofluorescence images of intestinal cryosections showing F4/80-positive macrophages (green), E-cadherin-positive epithelial cells (blue), and nuclei (DAPI, turquoise). Scale bars, 100 μm. **(B)** Relative abundance of macrophages in mice on normal water for 12 weeks (n = 3). Data are mean±s.d.

**Supplementary Fig 4.**
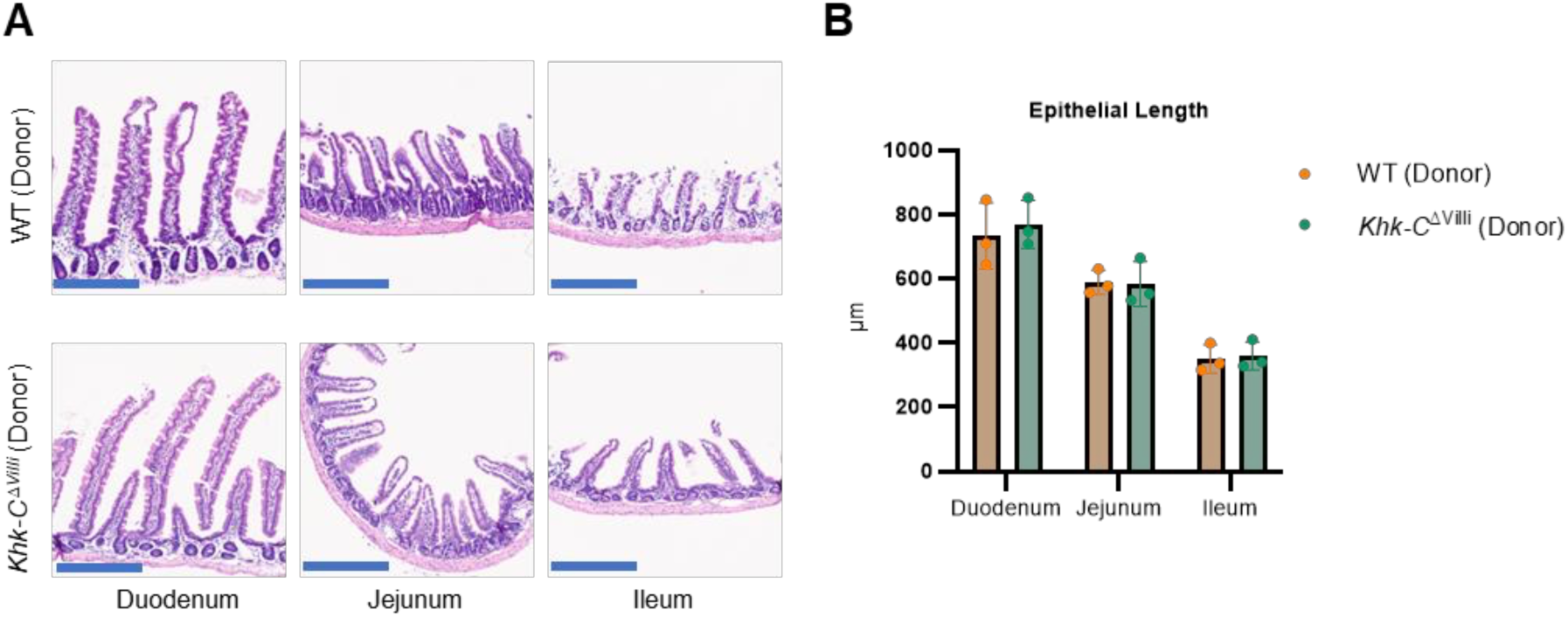
Fecal microbiota transplant does not alter epithelial length. **(A)** Representative H&E images of intestinal tissues in the recipient mice after 8 weeks of fecal transplant and HFCS drinking. Scale bars, 200 μm. **(B)** Epithelial length of intestinal tissues in the recipient mice after 8 weeks of fecal transplant and HFCS drinking (n = 3). Data are mean±s.d.

